# Visualizing genome synteny with xmatchview

**DOI:** 10.1101/238220

**Authors:** René L. Warren

## Abstract

In genomics research, the visual representation of DNA sequences is of prime importance. When displayed with additional information, or tracks, showing the position of annotated genes, alignments of sequence of interest, etc., these displays facilitate our understanding of genome and gene structure, and become powerful tools to assess the relationship between various sequence data. They can be used for troubleshooting sequence assemblies, in-depth sequence analysis, and eventually find their way in publications and oral presentations as they often translate complex and abundant data succinctly, with esthetically pleasing images. Here, I introduce xmatchview and xmatchview-conifer, two python applications for comparing genomes visually and assessing their synteny. Availability: https://github.com/warrenlr/xmatchview

---

In bioinformatics, daily use of ENSEMBL (https://www.ensembl.org) [1], the UCSC genome browser (https://genome.ucsc.edu) [2], and IGV [3] for visualizing genomics data is routine. The former allows for an easy-to-use visual navigation of the ENSEMBL genome databases. The latter two are customizable and flexible tools that can be used to situate sequence [read] alignments within a genome reference or draft assembly context, either online (UCSC) or as a stand-alone tool (IGV). These graphic interfaces are also useful to bioinformatics software development and debugging code as well as in *de novo* genome sequencing projects as they are incredibly effective for troubleshooting sequence assemblies. Circos, a highly cited stand-alone visualization tool, represents information and associated relationships as concentric circles and ribbons, allowing for abundant data (eg. human genome scale) to be depicted succinctly in full, within a computer screen window [4]. The success of circos is attributable in part due to its flexibility, versatility and customization in representing complicated relationships between data of all sorts, not just genomics. As attractive and convenient as circles are for displaying relationships between data, linear representations of synteny blocks between two DNA sequences remain intuitive. Here, I introduce xmatchview (https://github.com/warrenlr/xmatchview), a tool for visualizing DNA sequence alignments produced by cross_match (unpublished, http://www.phrap.org/), a robust implementation of the sensitive Smith-Waterman algorithm for DNA alignments. The software requires python and the python imaging library (PIL) to produce publication-ready images in a variety of formats (PNG, TIFF, GIF, and JPEG) and cross_match for performing the DNA alignments. With xmatchview, users can compare any two DNA sequences, including but not limited to gene reconstructions, genome assemblies, cDNA, nanopore reads, etc., and visually 1) identify collinear blocks, 2) assess the relationship between them, 3) analyze the sequence identity between repeated segments, and 4) view their frequency at given sequence coordinates.

As seen on Fig. 1, the xmatchview display consists of three main components: 1) The sequence objects, represented as black rectangles. Additional features such as exons, coding sequences, mRNA, ORFs, etc., are provided to xmatchview via a simple tab-separated file enumerating each start and end coordinates with, optionally, the color of each feature (plotted as yellow rectangles by default). Stretches of Ns in the reference and query sequences, when applicable, are shown in red on the sequence objects. 2) Relationships between co-linear block of sequences are represented by blue and pink polygons between the two black rectangles/sequence objects, depending on their direct or inverted associations, respectively. 3) A histogram on top of the reference sequence (upper most) black rectangle shows the sequence identity (top to bottom, from 0 to 100%) with the query sequence (lower rectangle). When a sequence is repeated, the color of the histogram changes to reflect its frequency. Visualizing repeat frequency is a feature unique to xmatchview that can be used to readily assess sequence complexity between (Fig. 1) or within (Fig. 2) sequences.

**Fig. 1.**
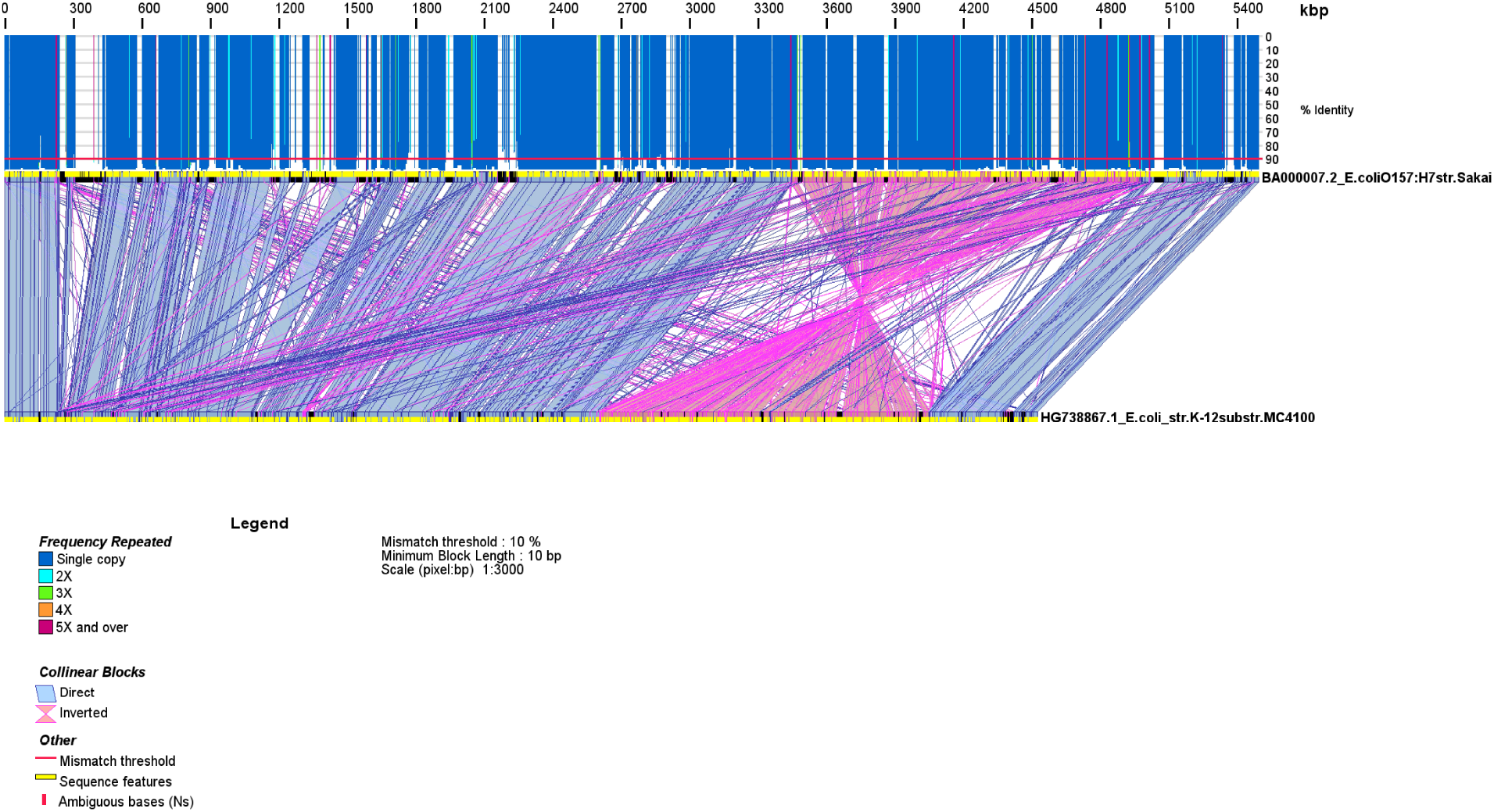
Genome sequence synteny between *E. coli* strains O157:H7 str. Sakai and K12 str. MC4100 (Genbank accessions BA000007.2 and HG738867.1). The genome sequence alignment was done with cross_match (options -minmatch 29 -minscore 59 -masklevel 101) and rendered with xmatchview (options -m 10 -r 10 -b 10 -c 2000 -a 200). The genomes of these two *E. coli* strains are largely co-linear, with the exception of a large inversion seen in one relative to the other. Although both strains comprise unique genomic sequences, strain O157:H7 has longer sequence stretches (up to 100 kb) not found in the K12 strain. Open reading frames in both genomes are displayed in yellow.

**Fig. 2.**
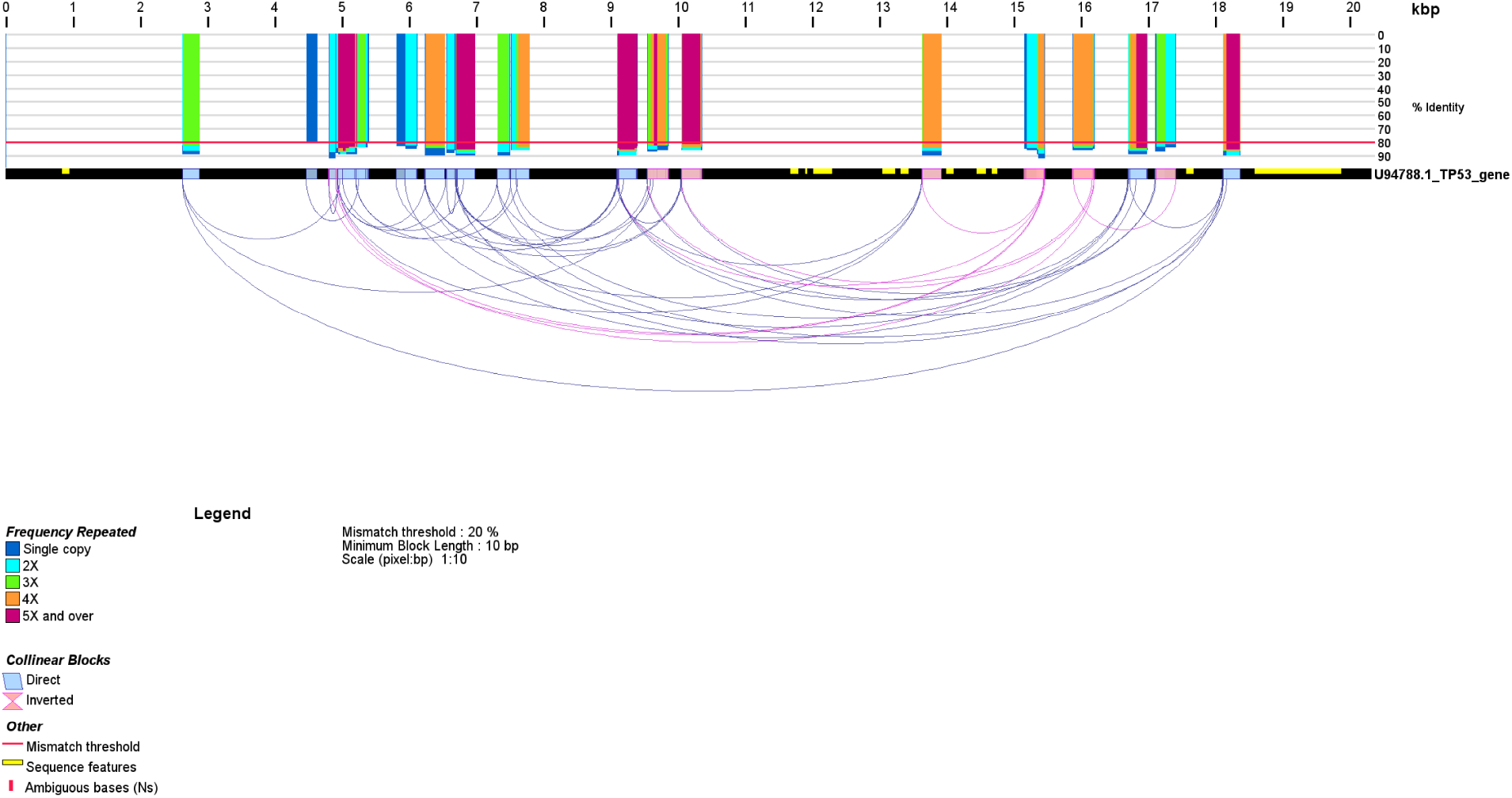
Sequence repeats within the human TP53 gene (Genbank accession U94788.1). The alignment was done with cross_match (options -minmatch 29 -minscore 59 -masklevel 101) and displayed with xmatchview (options -m 20 -r 10 -b 10 -c 10 -a 200). TP53 mRNA sequence coordinates within the gene are shown with yellow rectangles.

When the same sequence is given as input to xmatchview, internal repeats within that sequence are shown instead, representing only the reference sequence and the relationships between repeated blocks as arcs, instead of polygons (Fig. 2). Users can control whether to show the position of sequence features on the reference and query (-e and -y options), show co-linear blocks of a certain length (-b option) when their mismatch rates are below a threshold (-m option). The histogram is generated by moving a sliding window with a step length (-r recommended between 10-50). The color space in xmatchview is RGBA and the alpha channel is used for visualizing the relationship between co-linear blocks (-a option, transparent to solid, 0 to 255). Images from xmatchview have been used in a number of peer-reviewed publications to showcase genome co-linearity and/or highlight differences between DNA sequences [5] [6] [7] [8]. A modified version developed specifically for comparing conifer sequences with an evergreen tree representation, xmatchview-conifer, is co-released with xmatchview. The conifer tree representation differs from that of xmatchview. In the former, the sequence identity is color-coded within each synteny block relationship (light to dark shades of green tracking with increased sequence conservation), instead of a histogram (Fig. 3). Demo shell scripts that pipeline cross_match and xmatchview/xmatchview-conifer are included with the distribution and provided for guidance (runCompareTwoGenomesColinear.sh and runSpruceView.sh, respectively). Typically, sequences less than 10 Mbp in length are compared with cross_match and displayed in less than a few minutes using these pipelines, depending on your system. Both xmatchview and xmatchview-conifer are implemented in python and released under GPLv3.

**Fig. 3.**
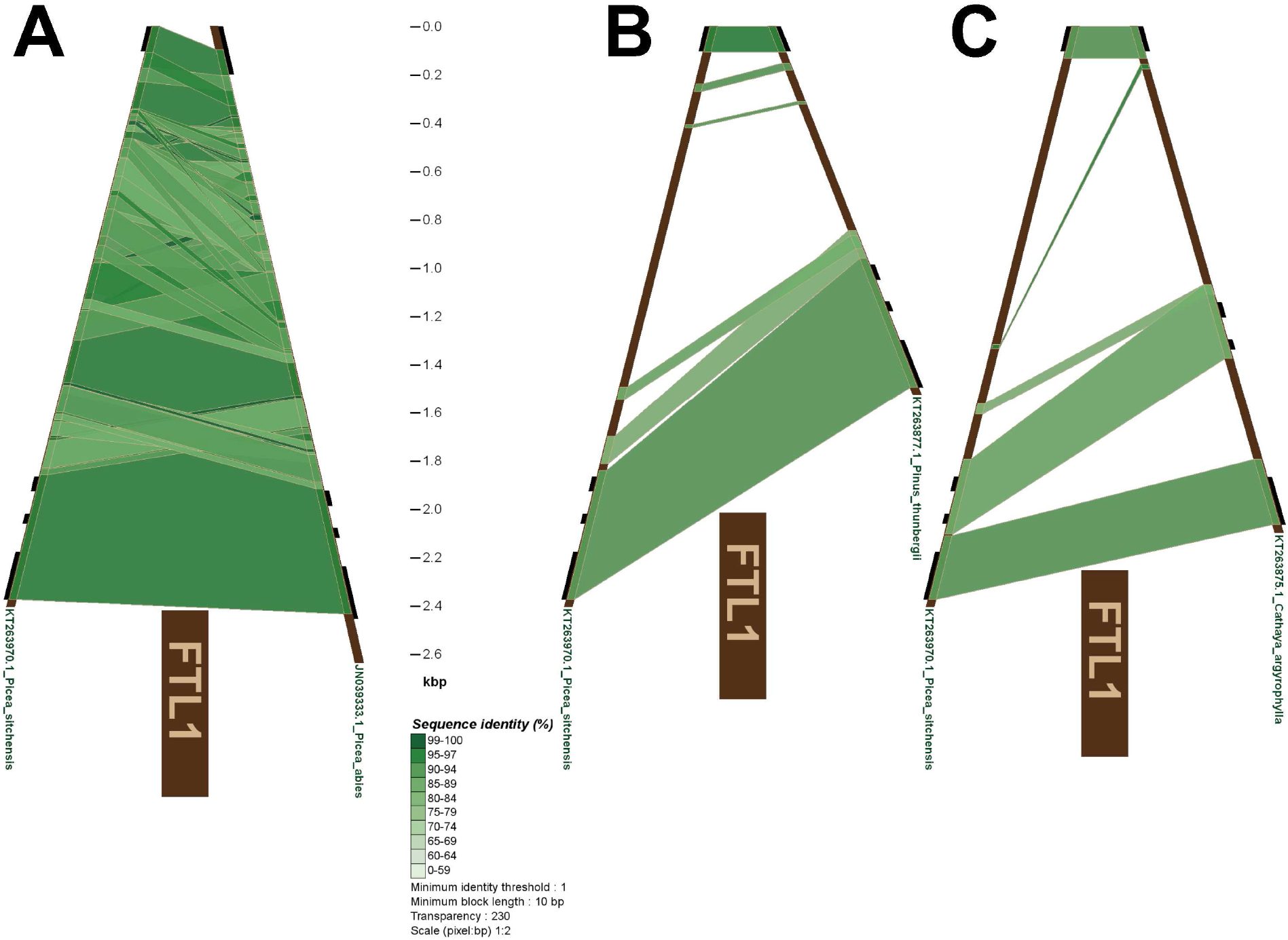
Sequence comparisons of the flowering locus gene FTL1 in selected conifer species of the order Pinales. The FTL1 gene for Sitka spruce (*Picea sitchensis*, Genbank accession KT263970.1) was compared to that of the A) Norway spruce (*Picea abies*, Genbank accession JN039333.1), B) Japanese black pine (*Pinus thunbergii*, Genbank accession KT263877.1) and C) Chinese mountain tree - yin shan (*Cathaya argyrophylla*, Genbank accession KT263875.1). The alignments were done with cross_match (options -minmatch 5 -minscore 10 -masklevel 101) and visualized with xmatchview-conifer (options -m 99 -b 10 -r 1 -c 2 -l FLT1 -a 200). The position of exons is indicated by the black rectangles outside on the outer edge of the tree representation. When comparing the FTL1 gene between the most distinct species (B and C comparisons), FTL1 sequence conservation (85-94% sequence identity) is mostly seen between exons.

## Acknowledgements

This work has been partly supported by the National Human Genome Research Institute of the National Institutes of Health (under award number R01HG007182). Additional funds were received through Genome Canada, Genome Quebec, Genome British Columbia and Genome Alberta for the Spruce-Up (243FOR) project (www.spruce-up.ca). The content reported here is solely the responsibility of the author, and does not necessarily represent the official views of the National Institutes of Health or other funding organizations.

